# Multi-omics alleviates the limitations of panel-sequencing for cancer drug response prediction

**DOI:** 10.1101/2022.06.15.496249

**Authors:** Artem Baranovskii, Irem B. Gunduz, Vedran Franke, Bora Uyar, Altuna Akalin

**Affiliations:** Non-coding RNAs and Mechanisms of Cytoplasmic Gene Regulation Lab, Berlin Institute for Medical Systems Biology, Max Delbrück Center (MDC) for Molecular Medicine, Hannoversche Str. 28, 10115, Berlin, Germany; Integrative Cellular Biology & Bioinformatics Lab, Saarland University, 66123 Saarbrücken, Germany; Bioinformatics and Omics Data Science Platform, The Berlin Institute for Medical Systems Biology, Max Delbrück Center (MDC) for Molecular Medicine, Hannoversche Str. 28, 10115 Berlin, Germany

## Abstract

Comprehensive genomic profiling using cancer gene panels has been shown to improve treatment options for a variety of cancer types. However, genomic aberrations detected via such gene panels don’t necessarily serve as strong predictors of drug sensitivity. In this study, using pharmacogenomics datasets of cell lines, patient-derived xenografts, and *ex-vivo* treated fresh tumor specimens, we demonstrate that utilizing the transcriptome on top of gene panel features substantially improves drug response prediction performance in cancer.

## Main

Cancer is a collection of diseases characterized by abnormal cellular growth and invasion of other body parts. It affected 19 million people in 2020 and was the cause of 9.5 m deaths that year alone ^1^. Cancer has been primarily considered to be a disease of the genome, where the accumulation of alterations in the genome is the underlying cause of how normal cells transform into malignant cancerous cells with improved survival and proliferation advantage ^2^. Such genetic alterations have been studied to understand the mechanisms of cancer and to develop therapies specifically targeted for such alterations. These targeted therapies and companion diagnostic tools have transformed oncology ^3^, promising more precise treatments tailored to tumors’ genetic profiles. Various targeted therapies have been successfully developed to counteract the defects in the molecular machinery borne out of such oncogenic mutations ^4–6^. To this date, most of the markers approved for targeted therapy decisions consist of single-gene markers ^7^. It has thus become crucial to develop accurate, sensitive, and high-throughput genomic assays to accommodate the increasingly genotype-based therapeutic approaches. Multiple approaches by commercial companies such as Foundation Medicine as well as large cancer research centers such as MSK and Dana Farber have produced their panel sequencing assays to guide therapy for cancer patients ^8^. These techniques examine frequently mutated genes in cancer to come up with mutations and copy-number variations for those genes. Especially for diagnostics, the approved methods for targeted drugs are usually the presence or absence of mutations. Therefore, the assay developers focus primarily on mutation calling accuracy as a metric of the usefulness and accuracy of the assay ^8^.

Although comprehensive genomic profiling using cancer gene panels has demonstrated value in broadening the treatment options for the patients based on matching a patient’s genomic lesions to cancer driver gene aberrations associated with FDA-approved treatment indications ^8,9^, the presence/absence of mutations in such genes does not necessarily translate into improved predictive power for estimating the patient’s response to the potential treatments. While for some drugs, the variation in drug response can be explained by a very specific mutation, for instance, BRAF V600E mutation is a strong predictor for response to Vemurafenib in metastatic melanoma ^4^, for many drugs, the knowledge of the mechanism of action is missing. This is because the majority of drugs are discovered via phenotypic screening of model systems rather than target-based approaches ^10^. Also, such single mutation markers for a given cancer type, are not necessarily good markers for other cancer types. For instance, BRAF V600E, while a good predictor for metastatic melanoma, is a poor predictor of response in metastatic colorectal cancer ^11^. More importantly, the latest compilation of the hallmarks of cancer recognized in the field includes not only genomic defects, but also factors such as the non-mutational epigenetic aberrations, involvement of the immune system in the tumor microenvironment, and the composition of the microbiome ^12^. Such layers of information that can be informative for how a tumor will respond to a certain treatment, cannot be fully captured by only focusing on the restricted set of genomic alterations, and necessitates other data modalities. Among those data modalities, transcriptome profiling –besides being a cheap and accessible option in terms of logistics– has been shown to yield strong predictors of drug response ^13–15^.

Here, we set out to quantify how much using the transcriptome as an additional data modality improves the drug response prediction performance compared to using only the genetic features restricted to the cancer gene panels (such as mutations and copy number variations). We leveraged publicly available pharmacogenomics datasets including genomic and transcriptomic profiles and drug sensitivity measurements in three types of datasets: cancer cell lines (using the CCLE database ^16^), patient-derived xenografts (PDX) ^17^ and *ex-vivo* treated fresh tumor specimens from Acute Myeloid Leukemia patients (BeatAML) ^18^. In all three settings, with an application of out-of-the-box machine learning techniques (*See Methods)*, we modelled drug responses for all available drugs in two reported data modalities, using only panel gene features (PS) or using the transcriptomic features on top of panel features (i.e. multi-omics (MO)). While achieving only moderate predictive power (mean R-square ∼ 10%, n = 396), the MO modality of the CCLE dataset showed an overall increase in predictive power (delta R-square ∼ 6%) in comparison to PS data (Figure 1a, Supplementary Table 1). Of note, we observed a significant positive correlation (*r* = 0.17, *p* = 0.0036) between the percentage of gene expression features among the top 100 features and an increase in MO’s predictive power over PS (Figure 1b). Modelling in xenografts generally conferred similar results. 4 out of 12 drugs showed a significant increase in MO’s predictive power of PS (Wilcoxon’s p < 0.05) (Figure 1c, Supplementary Table 2). While cancer cell lines and PDX models have been instrumental in developing cancer treatments, perhaps a better model for understanding the actual human drug responses would be the *ex-vivo* drug screening of fresh tumor specimens. Tyner *et al*. ^*18*^ have carried out a study of Acute Myeloid Leukemia tumor specimens, in which 106 drugs were tested on at least 100 samples each. These samples were also sequenced to profile both the genomic mutations and the gene expression levels. As we have observed in both CCLE and PDX datasets, we could reproduce our findings that using the transcriptomic features on top of mutations detected for cancer panel genes yielded a significant improvement in terms of the predictive power of drug response models (mean R-square of 11.8% for multi-omics and 8.2% for panel-seq; *p*<0.0001, one-sided Wilcoxon rank-sum test) (Figure 1d, Supplementary Table 3).

**Figure 1:**
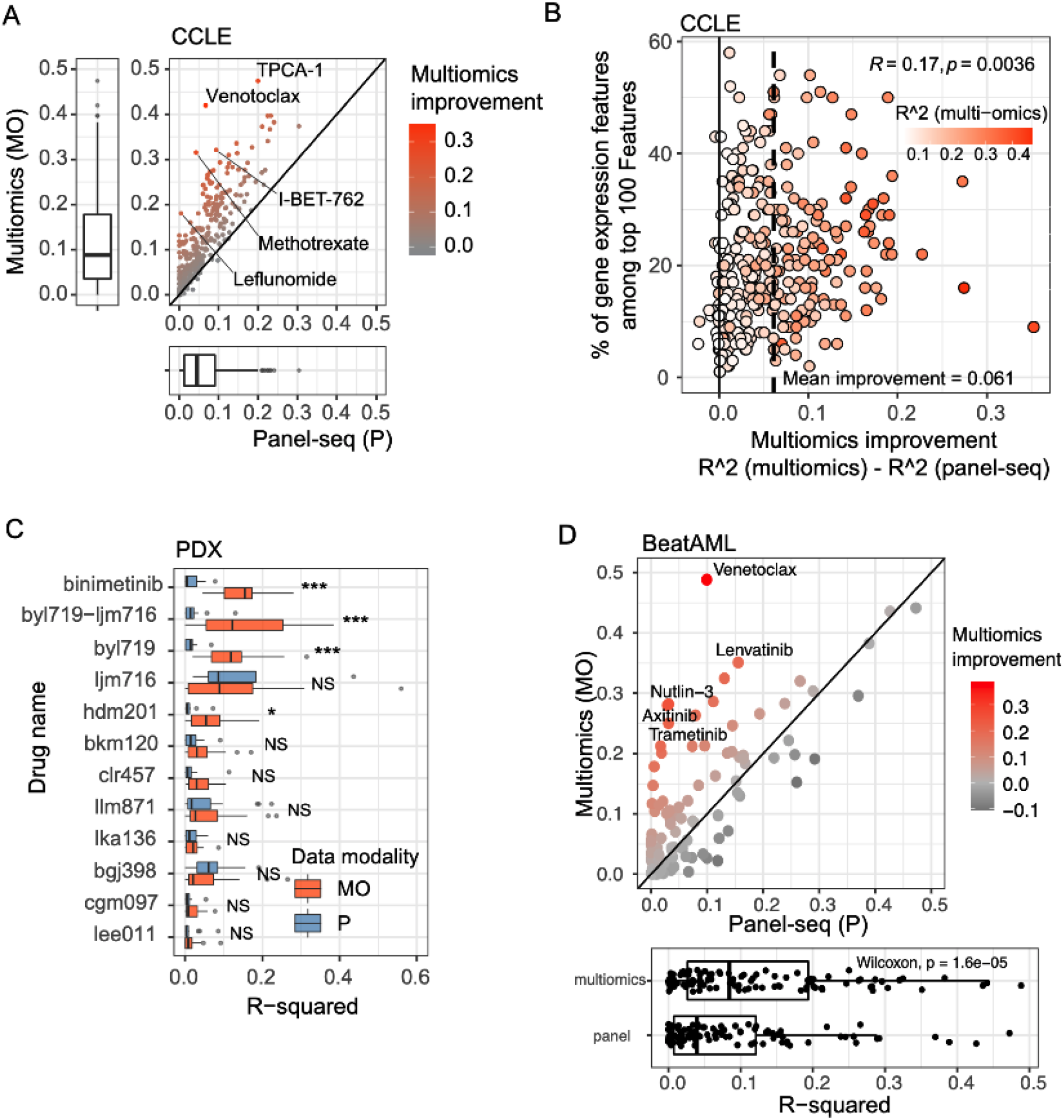
Evaluation of the performance of drug response prediction when using only panel-seq features (mutations and/or copy number variations) or using transcriptome features in combination with panel-seq features (multi-omics). A) Improvement of multi-omics (as in R-square metric) in comparison to panel-seq features for the testing portion of the CCLE dataset for 396 drugs. B) Correlation in the prediction performance improvement (multi-omics vs panel-seq) with respect to the proportion of the transcriptome features among the top 100 most important predictors of drug response for CCLE datasets. C) Improvement of multi-omics (in red) (as in R-square metric) in comparison to panel-seq features (in blue) for the testing portion of the PDX dataset for 12 drugs. D) Improvement of multi-omics (as in R-square metric) in comparison to panel-seq features for the testing portion of the BeatAML dataset for 106 drugs.

We believe that to better understand cancer and develop better drugs and diagnostics, we need to make use of all the molecular features by integrating different omic data sets. In this manuscript, using multi-omics and machine learning techniques, we show that multi-omics has indeed superior performance for drug response prediction in cancer.

## Methods

### Data/Code Availability

In this study, the following publicly available pharmacogenomics datasets were used:

- Cancer Cell Line Encyclopedia (CCLE) ^16^ downloaded from https://depmap.org/portal/download/
- Patient-Derived Xenografts (PDX) ^17^
- BeatAML: *ex vivo* drug sensitivity screening of Acute Myeloid Leukemia patient tumor specimens ^18^ downloaded and processed using the PharmacoGx R package ^19^.

All code to download, process, analyze the datasets and reproduce the figures in this manuscript can be found here: https://github.com/BIMSBbioinfo/multiomics_vs_panelseq

### Data Processing

#### Mutations

The mutation data were converted into a matrix of mutation counts per gene per sample. The resulting matrix was further filtered to only keep mutation data for genes in the OncoKB cancer gene list (https://www.oncokb.org/cancerGenes). Mutation data was available in all three datasets.

#### Copy Number Variations

The copy number variation data was used as downloaded from the respective resources. Copy number variation data was also filtered to only keep genes found in the OncoKB cancer gene list. Both CCLE and PDX datasets contained CNV data available, but it wasn’t available for the BeatAML dataset.

#### Gene Expression (Transcriptome)

To reduce the dimensionality and obtain less noisy features, the gene expression datasets were converted into gene-set activity scores using single-sample gene set scoring (singscore R package ^20^). The gene sets utilized in this study were the Cancer Hallmarks gene signatures (50 gene sets) from the MSIGDB database ^21^ and tumor microenvironment-related gene sets (64 gene sets) curated in the xCell R package ^22^. Gene expression data was available for all three datasets.

#### Drug sensitivity measures

For CCLE and PDX datasets, AUC (area under the curve) scores derived from dose-response curves as published in the respective resources were used in the prediction models. For the BeatAML dataset, recomputed AAC (area above the curve) scores were used as downloaded via the PharmacoGx R package ^19^.

### Drug response modelling with machine learning

For drug response modelling, we considered two main scenarios based on the availability of datasets. In the first setting, we considered mutation and/or copy number variation data for genes found in the OncoKB cancer gene list, which aims to simulate panel-sequencing. In the second set, we considered the features as in the first set along with the whole transcriptome profiling as an additional data modality, which is further converted into gene-set activity scores. The second set represents the multi-omics condition, in which the panel features are concatenated with the transcriptome features (gene-set scores).

For both settings, all three datasets (CCLE, PDX, BeatAML) were analyzed with the following protocol:

1. Only drugs that were treated on at least 100 samples were considered.
2. For each drug, the samples were split into training (70% of samples) and testing groups (30% of samples). See supplementary tables 1-3 for specific sample counts used for each drug.
3. Caret R package ^23^ was used to build random forest regression models (using either ranger ^24^ or glmnet ^25^ R packages) on the training data, where the genomic/transcriptomic features were used as predictors and the drug response values were used as the outcome variable. We used repeated 5-fold cross-validation for hyperparameter tuning to find the best model parameters based on the training data. Near-zero-variation filtering, scaling, and centring were applied as data processing steps. Applying PCA as a processing step led to poorer prediction results for both multi-omics and panel-seq features (Supplementary Figure 1a,b), therefore we excluded PCA processing. For the PDX samples, this step was repeated 20 times by resampling the training/testing portions. This was only applied for the PDX samples, due to the small number of drugs (N=12) treated on at least 100 samples.
4. The final model performance was evaluated on the testing data. Spearman rank correlation, R-square, and Root Mean Squared Error (RMSE) metrics were computed and the R-square metric was used to report the results in the final figures.

## Author Contributions

Data collection and formal analysis: A.B., I.B.G, V.F., B.U.; Writing and editing of the manuscript: all authors; Supervision: B.U., A.A.; Funding acquisition: A.A.

