## Supplemental Figure 1 for "Multi-omics alleviates the limitations of panel-sequencing for cancer drug response prediction"

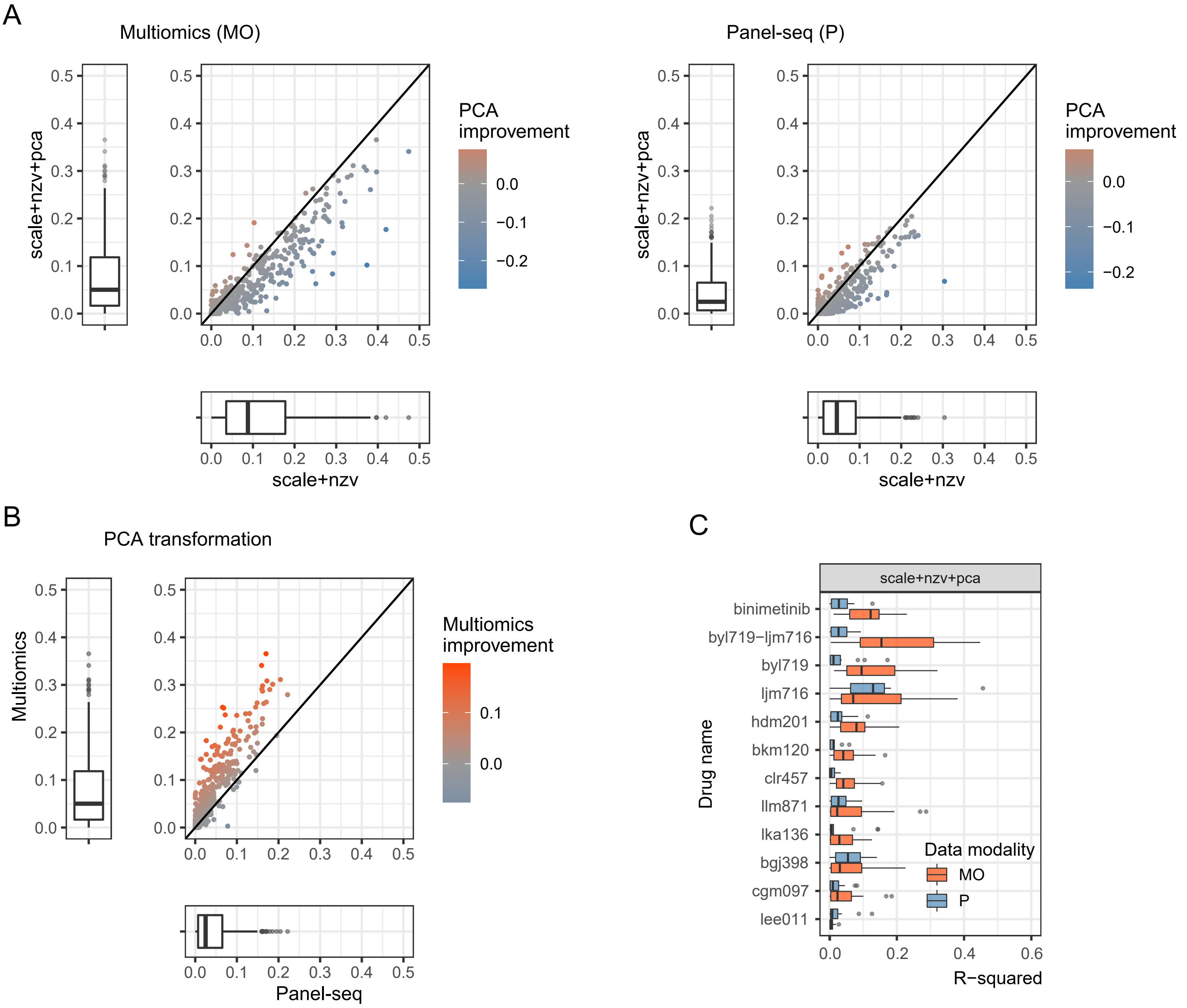


**Figure S1. Evaluation of PCA preprocessing's effect on drug response predictions.** A) Comparison of drug response predictions (as in R-squared metric) between scaled and PCA transformed (Y) and just scaled (X) estimated from *multiomics* (left panel) and *panel-seq* data for 396 drugs from CCLE dataset. B) Comparison of drug response predictions (as in R-square metric) between *multiomics* (Y) and *panel-seq* (X) PCA transformed data modalities for 396 drugs in CCLE dataset. Every point corresponds to a drug and is coloured by difference between *multiomics* and *panel-seq*. C) Improvement of *multiomics* (red) in comparison to *panel-seq* (blue) predictions (as in R-squared metric) estimated from the PCA transformed PDX dataset for 12 drugs.
